# Post-Ischemic Reorganization of Sensory Responses in Cerebral Cortex

**DOI:** 10.1101/2023.01.18.524583

**Authors:** P Hayley, C Tuchek, S Dalla, J Borrell, MD Murphy, RJ Nudo, DJ Guggenmos

## Abstract

Sensorimotor integration is critical for generating skilled, volitional movements. While stroke tends to impact motor function, there are also often associated sensory deficits that contribute to overall behavioral deficits. Because many of the cortico-cortical projections participating in the generation of volitional movement either target or pass-through primary motor cortex (in rats, caudal forelimb area; CFA), any damage to CFA can lead to a subsequent disruption in information flow. As a result, the loss of sensory feedback is thought to contribute to motor dysfunction even when sensory areas are spared from injury. Previous research has suggested that the restoration of sensorimotor integration through reorganization or de novo neuronal connections is important for restoring function. Our goal was to determine if there was crosstalk between sensorimotor cortical areas with recovery from a primary motor cortex injury. First, we investigated if peripheral sensory stimulation would evoke responses in the rostral forelimb area (RFA), a rodent homologue to premotor cortex. We then sought to identify whether intracortical microstimulation-evoked activity in RFA would reciprocally modify the sensory response. We used seven rats with an ischemic lesion of CFA. Four weeks after injury, the rats’ forepaw was mechanically stimulated under anesthesia and neural activity was recorded in the cortex. In a subset of trials, a small intracortical stimulation pulse was delivered in RFA either individually or paired with peripheral sensory stimulation. Our results point to post-ischemic connectivity between premotor and sensory cortex that may be related to functional recovery. Premotor recruitment during the sensory response was seen with a peak in spiking within RFA after the peripheral solenoid stimulation despite the damage to CFA. Furthermore, stimulation evoked activity in RFA modulated and disrupted the sensory response in sensory cortex, providing additional evidence for the transmission of premotor activity to sensory cortex and the sensitivity of sensory cortex to premotor cortex’s influence. The strength of the modulatory effect may be related to the extent of the injury and the subsequent reshaping of cortical connections in response to network disruption.

## 1 Introduction

The integration of sensory information within the motor cortex is an important feature of adaptive motor control. Sanes et al. [1] showed forelimb position modified motor cortex somatotopy, underlining the influence of sensory feedback on motor output. It is theorized that sensory prediction encodes an internal model necessary for generating task-specific forces which are then fine-tuned via prediction error when there is a mismatch between the projected trajectory and the resulting action [2]. While the cerebellum is generally thought of as the primary driver of error correction using ascending sensory information, recent evidence suggests that the somatosensory cortex also makes a significant contribution to this process [3, 4]. Photoinhibition of somatosensory cortex during a reach behavior limited adaptation to task perturbation, along with similar results from human somatosensory cortex inhibition during motor learning, add credence to the presence of sensorimotor interactions within the cerebral cortex that allow for adaptive motor behavior [5, 6]. Therefore, the reciprocal communication between somatosensory and motor cortex is likely an important part of the generation of complex volitional movements.

Disruption of sensorimotor connectivity, as occurs in acquired brain injury, often leads to a more severe motor impairment than what might be predicted based on damage to motor structures alone since there is an additional loss of sensory integration. In line with this, Shadmehr and Krakauer [7] propose that the deficits that arise with corticospinal tract injury are more than a downstream failure to transmit a control signal that may be expected with loss of corticospinal neurons. In their review, they draw parallels between subcortical stroke cases to patients with sensory dysfunction as they both display an inability to construct an accurate internal model for the movement. The loss of behavioral stability across trials in patients with motor impairments after stroke further supports the absence of integrated sensory predictions [8]. The basis for sensory dysfunction in motor impairment can be examined in greater detail in animal models with a focal cortical lesion [9]. Injury to motor cortex likely disrupts a wide array of neural connections, including those from sensory cortex [10]. The damage could directly affect axonal projections or indirectly cause the loss of input; in either case, the injury results in the loss of shared information between cortical areas [11, 12]. Even with some recovery via spared motor areas like premotor cortex and its descending motor projections, the loss of sensory input may still restrict full behavioral recovery. However, there is often novel, large-scale structural reorganization during spontaneous recovery to compensate for these losses and support the restitution of sensorimotor integration [13, 14, 15]. Thus, the post-injury plasticity of cortex is a potential substrate for re-establishing the sensory feedback that is necessary for motor strategies used in activities of daily living [16].

Somatosensory evoked potentials during motor learning are sensitive to task adaptation, leading us to consider the ways somatosensory responses within the cortex could be used as an index of sensorimotor function [17]. The somatosensory system codes information for both latency and magnitude of the response which are likely altered after injury as a part of ongoing plastic processes [18, 19, 20]. <u>The goal of this study was to uncover mutual communication between premotor and somatosensory cortex following recovery from a primary motor cortex lesion by measuring sensory related activity within the spared premotor cortex and modulation of sensory responses in somatosensory cortex by premotor activity.</u> Rats have a single putative premotor area known as the rostral forelimb area (RFA). While the majority of sensorimotor connectivity is mediated by the primary motor cortex (caudal forelimb area; CFA), direct, reciprocal corticocortical projections between RFA and somatosensory cortex (S1) also exist [21]. The anatomical connections and single premotor cortical area make rats a prime model for sensorimotor assays. If disruption of CFA and S1 connectivity through injury results in a subsequent restoration of sensorimotor integration through extant and de novo connections between RFA and S1, we expect that there will be modulation the cortical processing of the somatosensory response to peripheral stimulation.

To study this, rats were given a focal ischemic lesion within CFA. Four weeks later, the rats underwent a terminal procedure in which the neural responses to mechanical stimulation of the forepaw were recorded within the somatosensory and premotor cortex and modulated by intracortical microstimulation (ICMS) delivery. We found there are clear responses to peripheral somatosensory stimulation in both RFA and S1 after lesioning CFA that, while distinct, have a shared peak in spiking activity 50 ms after onset. This stands in contrast to previous work which resulted in the abolishment of a somatosensory response in premotor cortex without mediation of the primary motor cortex, reinforcing the idea that there is direct recruitment of premotor cortex with somatosensory processing during recovery [22]. Of note, rats with larger lesion volumes displayed a significant relationship for increased spiking during this later sensory response with a concurrent reduction in the early peak specific to S1 which could be a result of increasing sensorimotor integration between the cortical areas. The addition of a stimulus in RFA clearly disrupted patterns in the somatosensory response in RFA and S1 and predicted an increase in the weight of the shared response regardless of area. The modulation of the somatosensory response in S1 along with RFA by ICMS supports the establishment of their intercommunication in recovery. Together, these results provide evidence for a relationship between premotor and somatosensory cortex in sensory processing after primary motor cortex injury.

## 2 Materials and Methods

### 2.1 Animals

A total of 7 young-adult male Long-Evans rats (325g – 335g; 10 weeks on arrival; Charles River) were included in this study after meeting inclusion criteria on the behavioral task and surviving both surgical procedures (out of 12 animals). All procedures were approved by the Kansas University Medical Center Institutional Animal Care and Use Committee and complied with the *Guide for the Care and Use of Laboratory Animals (Eighth Edition, The National Academies Press, 2010)*.

### 2.2 Training procedures

Rats were initially trained to retrieve pellets from a semi-automated behavioral box [adapted from 23]. The task requires the rats to reach out of a slot in a Plexiglas behavioral box to retrieve a food pellet (45-mg Dustless Precision Rodent Pellets; Bio-Serv). The box is designed with a door which closes after each reach attempt through an infrared beam break, constituting a single trial. The trial was deemed successful if the rat grasped the pellet and brought it into the box without dropping it. Multiple initiations of reaches within a single trial were noted but were scored as successful if the retrieval was complete. The percentage of successful retrievals over all trials within a single session, which lasted for approximately an hour, was measured as an indicator of behavioral ability. Rats who reached 60 reaches within the first 15 minutes were stopped and the sessions were considered complete. Behavioral training took approximately two weeks, by which point rats were required to successfully retrieve pellets with a 60% success rate at least once prior to continuation in the study. By the end of the second week, 9 of 12 rats reached with sufficient motivation and accuracy to continue in the study. One of the original goals of these studies was to examine the effects of post-injury experience on sensorimotor integration. However, the differences between animals with rehabilitative training and those without were minimal compared to the variability in lesion size and corresponding functional impairments across groups (see Results), so the groups were collapsed for analysis.

### 2.3 Initial Surgery and Injury

Rats were anesthetized using an initial dose of isoflurane followed by bolus injections of ketamine i.p. and xylazine i.m. The animal’s anesthetic state was monitored and maintained using 0.1 cc intramuscular injections of ketamine (100 mg/mL), supplemented by xylazine (20 mg/mL) or isoflurane as appropriate. Six burr holes were made using a dental burr bit at 1, 0.5, -0.5 mm anterior/posterior and 1.5 and 2.5 mm lateral to bregma in order to target the caudal forelimb area (CFA) contralateral to the preferred reaching forelimb [24]. Three boluses of 110 nL of the vasoconstrictor endothelin-1 (ET-1) were delivered at each site for a total of 990 nL to create a focal ischemic lesion. Rats were administered buprenorphine and acetaminophen over the course of 48-hours following the injury procedure. In general, rats recovered well from the ischemic lesion surgeries.

### 2.4 Secondary Surgery and Data Collection

On post-operative day 26, all rats underwent a terminal procedure to test the ability to evoke and modulate the response to peripheral sensory stimulation in the forelimb somatosensory cortex (S1) and rostral forelimb area (RFA). After anesthetization and fixation in the stereotaxic frame, the incision was re-opened and a laminectomy was made at the base of the skull to allow drainage of cerebrospinal fluid and prevent edema. Two craniectomies were performed using a dental burr bit to expose the sensorimotor cortex of both hemispheres. RFA was identified through intracortical microstimulation (ICMS) mapping using standard procedures [25]. Using pulled glass micropipettes (3.5 M NaCl; platinum wire) with tapered tip diameters between 10-25-μm, motor areas were identified by lowering the microelectrode to a depth of 1500-μm using a microdrive (Narishige), delivering cathodal bursts of 13 stimulus pulses (1-Hz trains at 350-Hz), through a constant current stimulator (BAK) and ramping up current, up to 60 microamps, and observing any evoked movements. ICMS trains were coupled to an audio amplifier, enabling experimenters to associate joint movements with each train burst. The receptive fields of S1 were localized by inserting a microelectrode array (MEA) (NeuroNexus A4×8-5mm-100-400-703) into the target area connected to a digitizing headstage and acquisition hardware (Intan Technologies). The forelimb area of S1 was defined by the ability to evoke output by manually palpating the contralateral paw while listening for audible spiking responses on different sites within the array. The first insertion site with any responsive channels was used to avoid exacerbating edema with multiple insertions of the MEA. A more detailed topography of forelimb somatosensory cortex was then resolved by switching to Von Frey microfilaments. Following identification of S1 and RFA, two MEAs were lowered into these areas using micropositioners. Response properties of neural units were recorded from 4-shank (shank-spacing of 400-μm) polymeric MEAs with 8 electrodes (site-area of 703-μm^2^) per shank (site-spacing of 100-μm; NeuroNexus A4×8-5mm-100-400-703) with a sampling frequency of 30-kHz (Intan Stimulation/Recording Controller). Additional application of isoflurane was critical to ensure that large deflections in the local field potential that appeared to be related to whisking were mitigated and did not contaminate the recorded response to sensory stimulation. Stimulation was delivered using two modalities: electrical microstimulation delivered in the premotor cortex and mechanical sensory stimulation of the forelimb using a solenoid. Three types of trials were cycled through during the neural recordings: solenoid-only peripheral stimulation (“Solenoid”), intracortical microstimulation-only (“ICMS”), or both types of stimulation at a set latency for each experiment (“ICMS + Solenoid”). A miniature 5V solenoid (ID 2776; Adafruit) was clamped to the stereotaxic apparatus and positioned next to the forelimb to contact a site which strongly elicited an increase in spiking on the MEA in S1. The solenoid was placed at a distance where the small pin contacted the forepaw during its range of movement and displaced it with a load less than 80g. The solenoid was activated in 1-Hz intervals. While the solenoid is not as sensitive as force-controlled mechanisms like that described by Emanuel et al. 2021, it is similar to the setup described in Foffani et al. [26] and Moxon et al. [19]. For the ICMS trials, a single cathodal-leading, bi-phasic square wave stimulation pulse with phase duration of 100-μsec per phase was delivered using a single electrode site from within the recording array in RFA. A digital signal generator (Master-9; A.M.P.I.) was used to cycle through logical combinations of digital inputs to the combined recording and stimulation unit so that each experiment cycled serially through 100 trials each of the three types of trials. A stimulation-state machine module embedded in the FPGA of the recording and stimulation unit was used to implement any delay between solenoid and ICMS, or between ICMS and the onset of the trial indicator (Figure 1).

**Figure 1.**
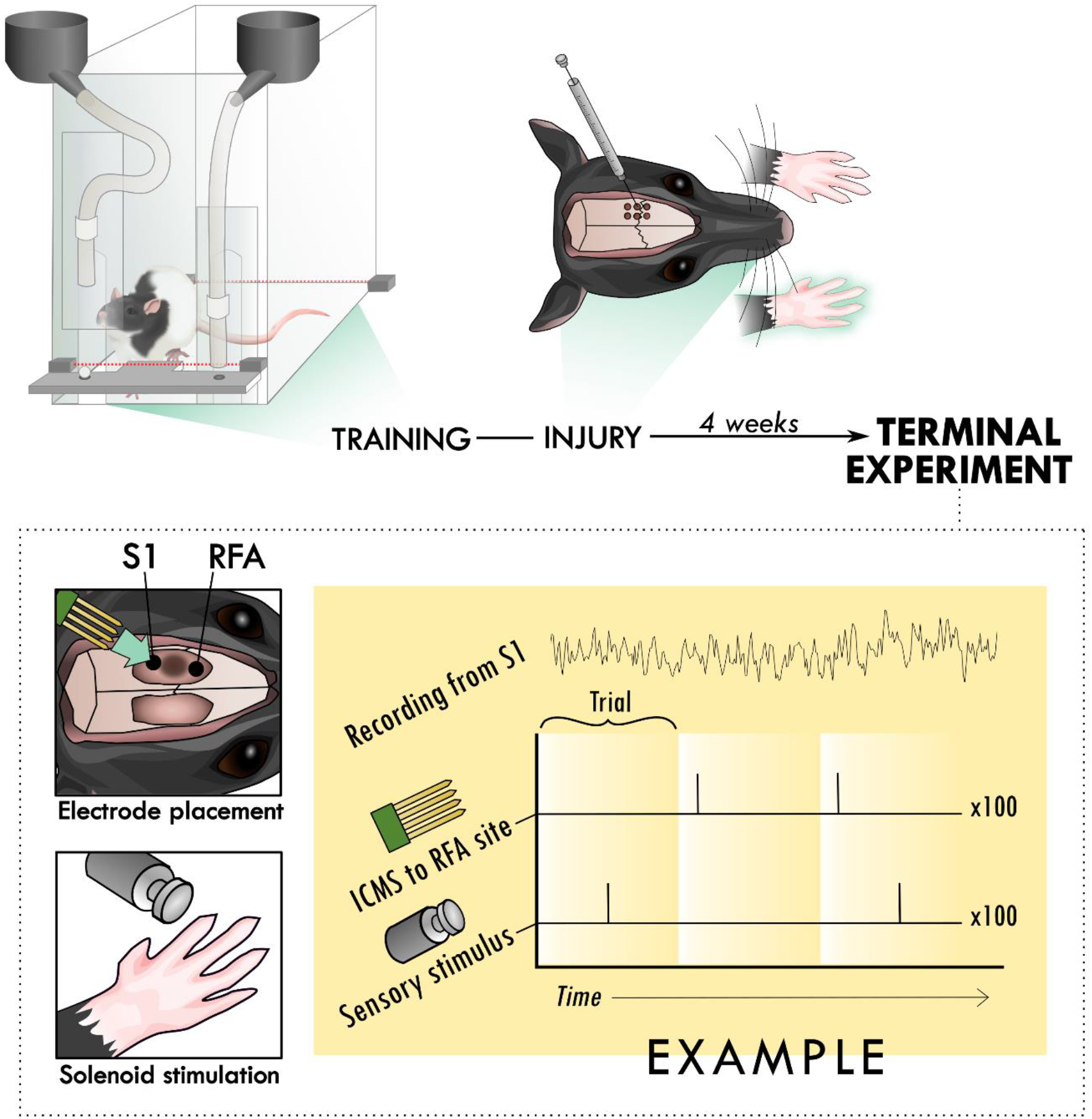
Experimental set-up and design. Rats were trained to perform the skilled reach task using an automated behavioral box. After training, rats underwent an injury procedure in which 6 burr holes were made over primary motor cortex and used to inject the vasoconstrictor, endothelin-1. The resulting lesion affected the contralateral forelimb as assayed in the skilled reach task. 4 weeks after the injury procedure, a terminal procedure was conducted in which bilateral cranial openings were made to expose cortex. Two microelectrode arrays were placed in putative premotor (RFA) and somatosensory cortex (S1), near the lesion in primary cortex and the solenoid was positioned to deliver peripheral sensory stimulation. Trials were cycled between peripheral stimulation only, intracortical microstimulation (ICMS) only, and the two together at an offset latency while recording from both arrays.

### 2.5 Histological Preparation

Immediately following the terminal procedure, the rats were injected with (Beuthanasia-D) and transcardially perfused with 0.1M saline solution with heparin and lidocaine followed by 4% paraformaldehyde. The brains were extracted, gelatin embedded, and sectioned coronally at a thickness of 50-µm using a freezing, sliding microtome. The sections were mounted on slides and stained with cresyl violet. An example of coronal hemisections corresponding to CFA are shown with examples of the small, subtotal lesions generated in these experiments is shown in Figure 2. Sections were analyzed under a light microscope (Zeiss AxioImager M2) and sampled at 1.2-mm intervals. Hemispheric volume through the sensorimotor cortex was estimated using the Cavalieri method, labelling the region of interest, which was drawn around gaps and non-viable tissue in the injured cortex to outline the lesion area, with markers (Figure 2B).

**Figure 2.**
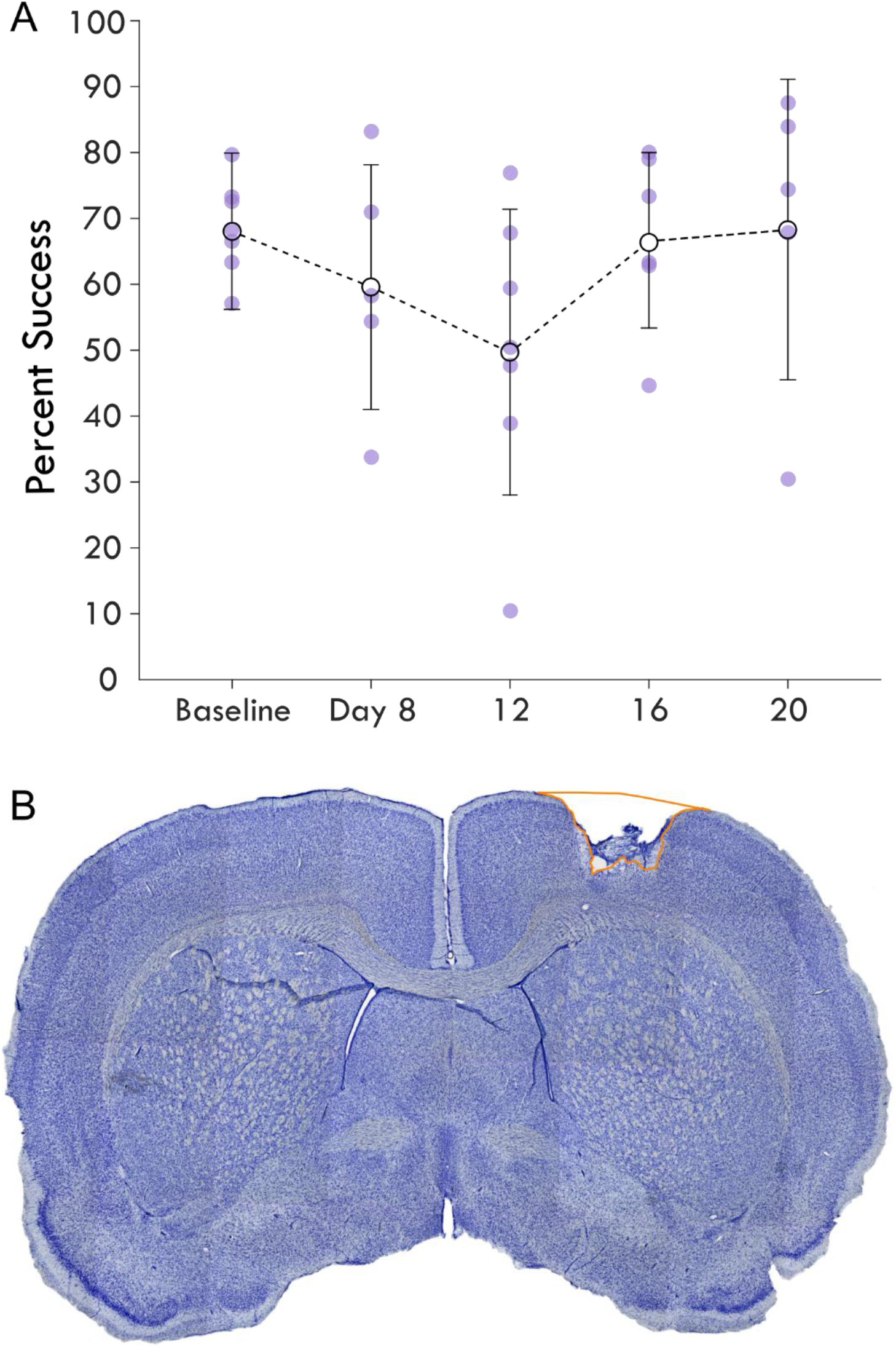
Average single pellet reach task success following cortical lesion. Panel A shows the percent success or the percent of successful retrievals out of total attempts made in each assay is shown over the experimental course. The baseline behavioral score averages pre-injury assays while the next timepoints represent assays post-injury. Each rat’s success is plotted as a point and shown with the mean and standard deviation at the timepoint for each assay. Panel B highlights a coronal section of a brain stained with cresyl violet showing an example cortical lesion.

### 2.6 Neural Data Processing

#### 2.6.1 Spike Detection

Unit activity was filtered using a 4th-order elliptic IIR filter with passband cutoffs of 300-Hz and 5,000-Hz. After bandpass filtering, a virtual common reference was estimated as the ensemble mean of all channels on a given MEA, which was then subtracted from each individual channel signal. After filtering, spikes were detected using a smoothed nonlinear energy operator (SNEO) [27], with a smoothing width of 5 samples and a minimum amplitude threshold of 15-μV. This detection method captured the physiological response of multi-unit activity in a manner that was consistent with previously reported results [28, 29], indicating its acceptability as a proxy for neural excitation and inhibition in response to the peripheral sensory stimulus. For all trials, any absolute deviation greater than 450 µV was considered artifact and spikes 4 ms before or after the artifact were discarded.

#### 2.6.2. Spike Unit Analysis

The spike rate was determined by taking the square root of multi-unit spike counts in 5 ms bins. Mean baseline spike rates were computed on a per-channel basis, averaging together the 10 time-samples beginning 50 ms prior to the first stimulus of each trial, yielding a time-series of 10 mean square-root-transformed spike counts for each channel. These values were then used on a per-channel basis to compute both a mean and standard deviation; the threshold for significant evoked activity was counted as this mean value plus 3 times the standard deviation computed in this way. Channels from a given trial with a baseline spike rate below 2.4 √spikes/s (< 1% of all data collected) were removed from consideration because the low event rate would be insufficient to power statistical analyses. For each peri-event time histogram, the square-root-transformed spike counts were computed in the same way for each grouped experimental variable (i.e. trial type and area) and plotted for each bin. A Savitzky-Golay Filter was applied to smooth the spike rates to find the corresponding 95% confidence band at each time point. Smoothing used a 21-sample Kaiser window with shape parameter set to 38 to fit a 3rd order polynomial to the sequence of spike times.

### 2.7 Component Analysis

The spike rates of channels for each recording block were analyzed by trial type to determine their top-3 principal components, collapsing both animal and area in the process. Using the principal component coefficients and the mean rates, the components were reconstructed to maximize independence from each other using reconstructed independent component analysis (r-ICA). Both principal and independent component coefficients were plotted over trial time and compared for fit to data as described by the percent explained. Each component is an eigenvector on which the average spike rates of channels were plotted by their respective component weightings. The component weights capture the strength of each component and its features in the average spike rate of a channel and were used to reconstruct the original rate by multiplying the weights for each component by the independent component coefficients. Finally, the mean independent component weights by area and trial type were averaged as a measure of change in any features of the spike rate. A built-in algorithm supplied by MATLAB (R2017a+) was used for r-ICA which has an in-depth description within the MATLAB documentation, but is summarized here for reproducibility: The r-ICA algorithm maps input data to output features by minimizing a standard limited memory. In this case, we let each observation *x* be the set of each mean spike rates for a given channel; we would like to recover both *s* and *A* such that each column of *s* is statistically independent from the other while still allowing us to accurately reconstruct each *x* in the observed data:

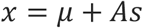

- *x* is a column vector of length p.
- *μ* is a column vector of length p representing a constant term.
- *s* is a column vector of length q whose elements are zero mean, unit variance random variables that are statistically independent of each other.
- *A* is a mixing matrix of size p-by-q.

To provide a way to fit this regression, we require that the initial weights for *A* are set to the top-*k* principal components for *X*, the set of all mean spike rates (such that rows of *X* are time samples and each column of *x* is a different channel’s spike rate). In these data, we selected *k = 3*, as the top-3 principal components explained most of the observations.

### 2.8 Statistical Models

General Linear Mixed Effect models were constructed to test the predictive power of different factors in the weights of the independent components. To fit models, we used the MATLAB (R2020a) ‘fitglme‘ function to fit data with a binomial distribution and logit link function. All models were fit with the ‘DummyVarCoding’ parameter set to ‘effects’ (the coefficients must sum to zero), and ‘FitMethod’ parameter set to ‘REMPL’ (restricted maximum pseudo-likelihood estimation for the model optimizer objective function). Models only included the Solenoid and ICMS + Solenoid trials to primarily determine the effect of a preceding ICMS pulse on the Solenoid response. Two models were created to assess Components 2 and 3 and included terms for the intercept as well as area, lesion volume, trial type, and all their interactions.

### 3 Results

To investigate the influence of injury on the connection between sensory and premotor cortex, rats had an ischemic lesion induced in the forelimb area of CFA (Figure 1). The lesion was more variable across rats than previously described, with a mean volume of 5.4 ± 3.7 mm^3^ [24, 30]. The rats were assessed in the skilled reach task to determine severity of motor deficits and recovery profiles. Figure 2A shows the rats’ behavioral scores on Days 8, 12, 16 and 20 after the ischemic lesion. As expected, average performance on the reaching task dropped after injury and recovered towards pre-lesion levels by the time the stimulation assays were performed. The majority of the rats had at least one timepoint where their behavioral performance was significantly reduced compared to baseline (Table 1); however, there was a large amount of variance in reaching success overall. Injection of ET-1 was done to target all cortical layers of CFA; however, the resulting lesion had a range of presentations. Categorizing lesions by their anatomical extent explained most of the variance in behavioral performance but did not result in clear patterns in spiking activity (Supplementary Fig. 2). An example lesion is shown in Fig. 2B.

**Table 1.** Difference of each rat from baseline behavioral scores at each timepoint. Standardized scores relative to a two-tailed T-distribution with a p value of 0.01 (t = 2.898, d.f. = 17) are highlighted (yellow values are below the confidence interval and gray those above). Timepoints with a dash represent days where the rat would not attempt reaches while N/A shows timepoints where the assay data is unavailable.

| Animal | Lesion Volume (mm<sup>3</sup>) | Standardized Scores |  |  |  |
| --- | --- | --- | --- | --- | --- |
|  |  | Day 8 | Day 12 | Day 16 | Day 20 |
| MM-W1 | 1.4 | -- | -3.2740 | 4.0961 | N/A |
| MM-U1 | 1.7 | 0.8578 | -0.2716 | 1.7121 | 5.4936 |
| MM-S1 | 3.1 | 5.2291 | 2.9809 | 3.7315 | 6.7804 |
| MM-V1 | 4.8 | -5.0754 | -7.4630 | -2.0480 | N/A |
| MM-T2 | 6.5 | -12.438 | -10.605 | -8.5496 | -0.2645 |
| MM-U2 | 8.8 | -3.6815 | -6.4801 | -1.8836 | 2.0838 |
| MM-T1 | 11.2 | -- | -20.777 | -- | -13.629 |

Four weeks after the injury procedure, rats underwent a terminal procedure in which the response to peripheral somatosensory stimulation was recorded in RFA and S1. In a subset of trials, ICMS was delivered in RFA alone or preceding the solenoid stimulation. Figure 3 shows the averaged responses of each trial type in both cortical areas. A peak in spiking activity was visible in both RFA and S1 following solenoid-only stimulation. In S1, this response was bimodal with an early increase followed by a second peak 50ms after solenoid onset; while in RFA, there was a peak which was similar in profile and locked to the same latency as the later peak in S1. With ICMS-only stimulation, there was a peak in spiking activity within RFA immediately following the pulse that was not observed in S1. In the ICMS + Solenoid condition, there was no obvious modulation in the profiles of the somatosensory-evoked peaks in either area. None of the averaged responses to any of the trial types exceeded the threshold of 3 standard deviations above baseline firing; however, the shaded confidence intervals did in some of the cases as highlighted in the inset of Figure 3. The significant variability suggests that there may be differences in the response at a channel level, leading to the later breakdown of the channel averaged activity by their main features.

**Figure 3.**
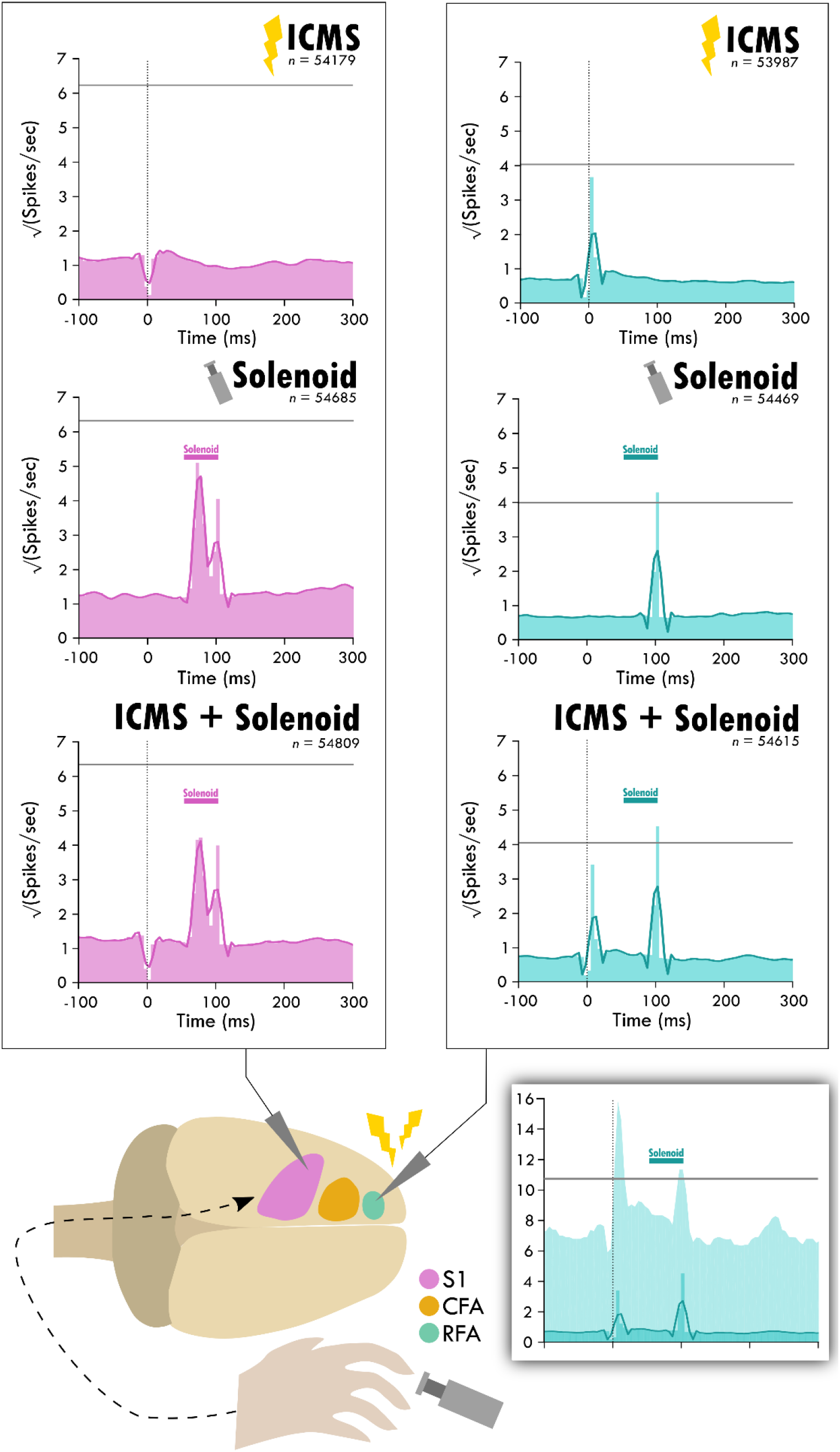
Average response in spiking activity to stimulation type in sensory and premotor cortex. The schematic at the bottom shows the cortical areas and the respective recorded activity during each of the three trial types: ICMS, Solenoid, and ICMS + Solenoid. Each peri-event time histogram shows the averaged spike rate for a trial type in each respective cortical area. The lighter shaded bins show the mean spiking at each time point while the overlaying dark line is the smoothed mean. The horizontal gray line is one standard deviation above the average pre-stimulus activity. The vertical dotted line shows where ICMS delivery occurs; spikes 4 ms around any artifact, as occurs with ICMS delivery, were removed from consideration. The shaded gray area shows the extent of solenoid action from the trigger to its retraction time. The shaded inset on the bottom shows the PETH plot of ICMS + Solenoid trials in RFA with the 95% confidence interval of the smoothed mean shown as a lighter background and a horizontal gray line for 3 standard deviations above the average pre-stimulus activity.

The average spiking by trial type was used to identify the principal components of the response to peripheral sensory stimulation from which the independent components were derived. The top independent components help determine underlying trends through dimensionality reduction, explaining a large portion of the data (Supplementary Figure 3,4). Figure 4B shows an example of the contribution of the top-3 independent components to the somatosensory response with their respective weightings. The top-3 components for each trial type are displayed in Figure 4A starting at 50ms after the zero point when sensory stimulation begins. The components for the solenoid only and ICMS + Solenoid trials are similar to one another with a relatively flat first component while the next two components have peaks in values at 50 and 100ms respectively. The combined ICMS + Solenoid trials has an additional negative dip in the coefficient of Component 2 at 100ms that is not seen in the Solenoid-only trials. In comparison, the ICMS-only trial components have a different pattern than the other trial types. Component 1 is more time invariant whereas Component 2 has a strong positive relationship to the neural data at the trial outset that decreases over the course of the trial to become negative. Component 3 features peaks at 70ms and 260ms; however, between the two peaks the coefficient drops to becomes negative. Underneath the representation of the independent components, the clustering of the average spike rate for each individual channel in a single experimental block is plotted along the new axes based on their weightings. Overall, the scatterplots highlight patterns in the spiking response along the range of independent features. The insets further capture the descriptive power of the combination of component weightings by displaying the mean spike rate for exemplar points along the component axes. The independent features and their respective weightings can be used to reconstruct individual trial rates, capturing the main trends (Fig. 4C). For the rest of the analysis, Component 1 was left out as it represents a trial invariant response to stimulation.

**Figure 4.**
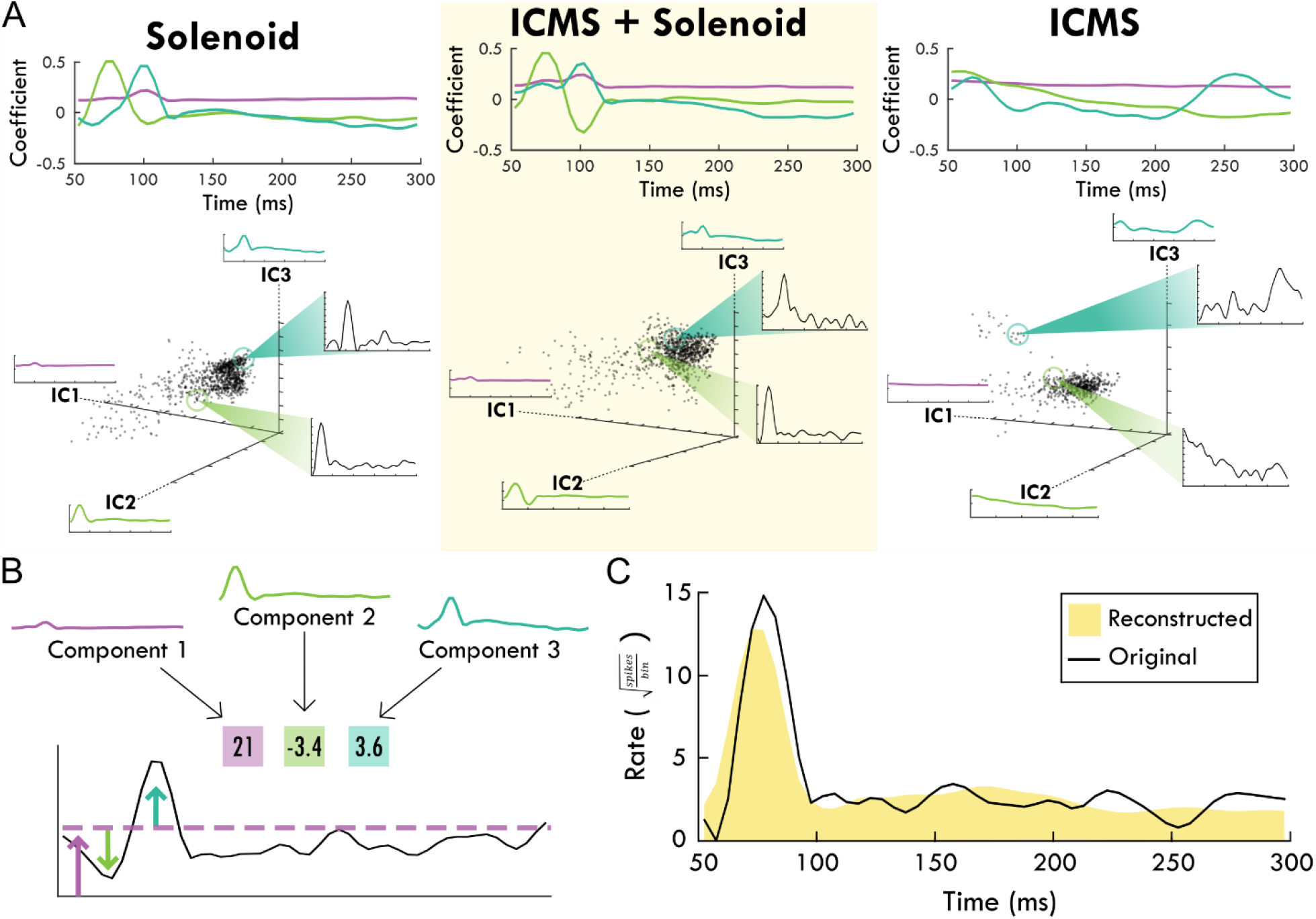
The top-3 independent components of the spiking response of each trial type. Panel A shows each stimulation type and the corresponding independent components over the trial time. Positive coefficients are correlated with spiking activity while negative coefficients are anti-correlated with spiking activity. In the scatter plots below, each component is shown as an axis and each trial is plotted as a point within the three dimensions. Exemplar trials are highlighted and shown in insets with spike rate over time. Panel B shows how the component weights (boxes) scale the component shapes to describe the features of the mean firing rate of an example channel. The corresponding blue and green arrows point to the deviations in mean firing rate while the purple arrow and line generally indicate the background firing rate that are captured by the respective component and its weight. Panel C shows the reconstruction (shaded yellow) of the mean spike rate of an example channel (black line) using the descriptive weightings of the independent components.

Weights for trials were plotted on the axes for Components 2 and 3 to examine trends in the response between Solenoid and ICMS + Solenoid trials by cortical area with breakout panels showing the spike rate of individual trials (Fig. 5). The weights along Components 2 and 3 independently capture the early and late portion of the sensory response to represent the peaks in the channel averaged activity (Fig. 5A and a,b). In the Solenoid trials, there is a single cluster for RFA that varies primarily along Component 3. Trials in S1 have a similar cluster with features that resemble that of RFA (Figs. 5A and c,d); however, there is also an additional smear of trials that extend out along the Component 2 axis that have an early increase in spike firing independent of a later peak (Figs. 5A and a,b). With ICMS + Solenoid trials, the patterns of weights shift to be diffusely distributed along both axes (Fig. 5A). The combined ICMS + Solenoid stimulation seems to modulate the spike rate in both areas on an individual channel basis, shifting the peaks both positively and negatively (Fig. 5e,f). The change in the sensory response with a preceding ICMS pulse is exemplified in Fig. 5B where the weights of the channel averaged activity are plotted for Solenoid and ICMS + Solenoid trials in two animals. In both, there is a clear shape to the shift in weights between trial types.

**Figure 5.**
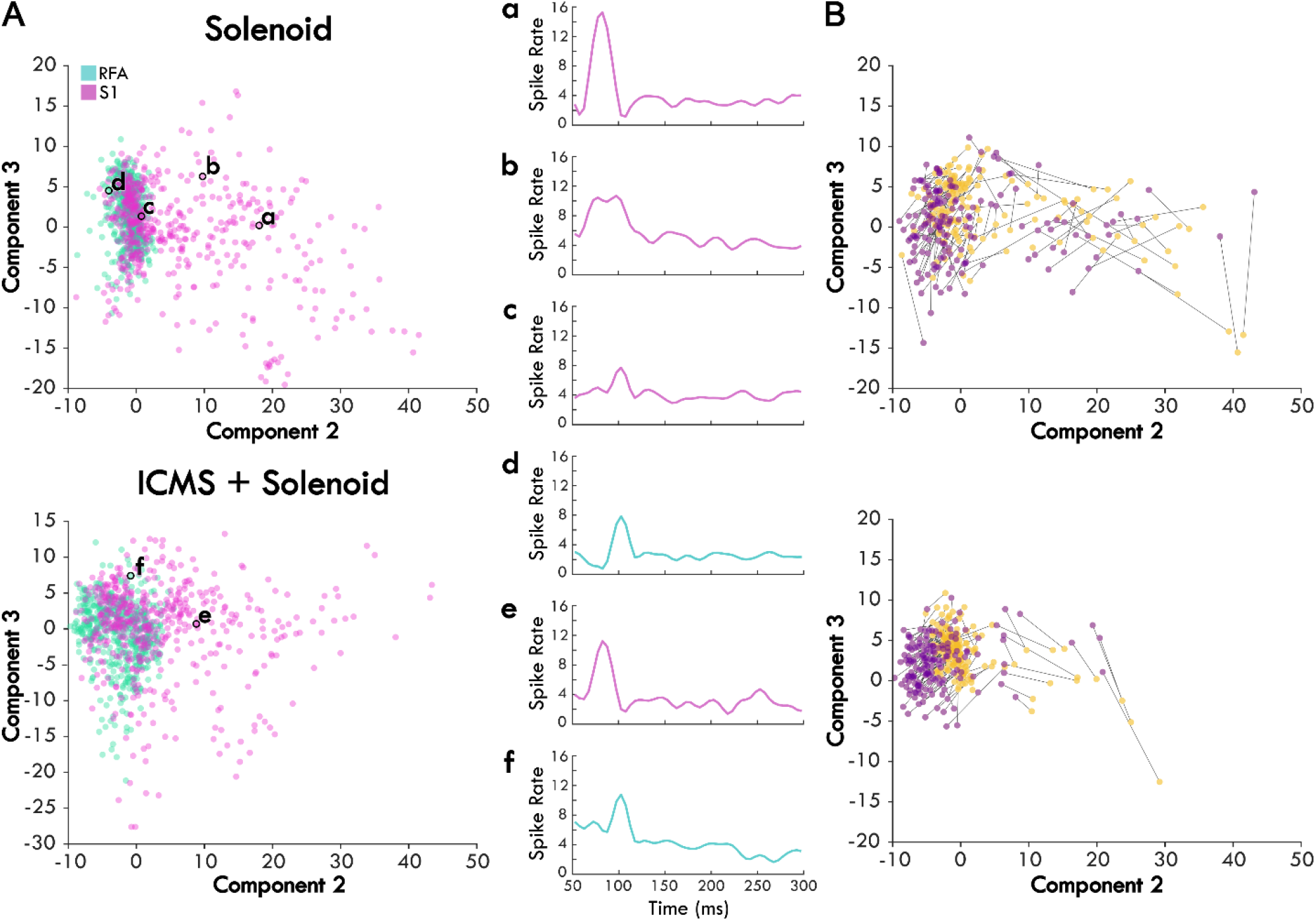
The average scores of Solenoid and ICMS + Solenoid trials in each area. Panel A displays each channel’s spike rate averaged across an experimental block and plotted on the second and third component axes based on its score. Channels in RFA are shown as blue and those in S1 are pink. Example channels are outlined in black and shown in the subpanels. a and b. two different channels within S1 of the same rat showing different response profiles. c. channel in S1 of the rat with the largest lesion volume highlighting a smaller first peak. d. channel in RFA showing a single evoked peak. e. and f. the same channels in panels a and d respectively during ICMS + Solenoid trials. Panel B shows the change in individual channel component scores without reference to cortical area between Solenoid (yellow) and ICMS + Solenoid trials (purple) in two exemplar rats.

To consider the difference of area on the response to stimulation, the weight of each component was averaged independently for RFA and S1 (Fig. 6A). The mean weights for Component 1 are comparable with little variance and are similar across trial type while the mean weights of Components 2 and 3 are differentially affected by the area and stimulation type. To quantify these differences, a general linear mixed model was created to determine the significance of the different variables on the response. The model only considered the stimulation types Solenoid and ICMS + Solenoid since we wanted determine if the ICMS stimulation modulated the sensory response to a solenoid strike. Area has a significant effect on Component 2— S1 is predicted to produce greater mean values than those in RFA (RFA = -3.9527± 0.29291, d.f.= 1946, p<0.0001). This is represented in Figure 6 where Component 2 overall has a positive weight in S1 and a negative weight in RFA in both Solenoid and ICMS + Solenoid trials. The negative weight of Component 2 in RFA means that the feature shown in Figure 4 is inversely present in the data. Additionally, the interaction of area and lesion volume on the mean weight of Component 2 is also significant where larger lesion volumes predict more positive values in RFA and more negative values in S1 (RFA = 0.28152± 0.044376, d.f.= 1946, p<0.0001). There is no effect of area or lesion volume alone on Component 3; however, area and lesion volume together have a significant interaction. The mean of the component is predicted to become more positive in S1 as lesion volume increases whereas it becomes more negative in RFA (RFA = -0.17202± 0.031232, d.f.= 1946, p<0.0001). Stimulation type is a significant predictor for differences in the mean of Component 3 and not Component 2 (Solenoid = -0.8183 ± 0.17744, d.f.= 1946, p<0.0001; Solenoid = -0.092543 ± 0.25236, d.f.= 1946, p=0.71388). Component 3 is weakly present on average in both S1 and RFA Solenoid trials (Fig. 6A). However, with the addition of ICMS in the ICMS + Solenoid trials, the weight of the component increases in both areas with Component 3 featuring stronger in S1.

**Figure 6.**
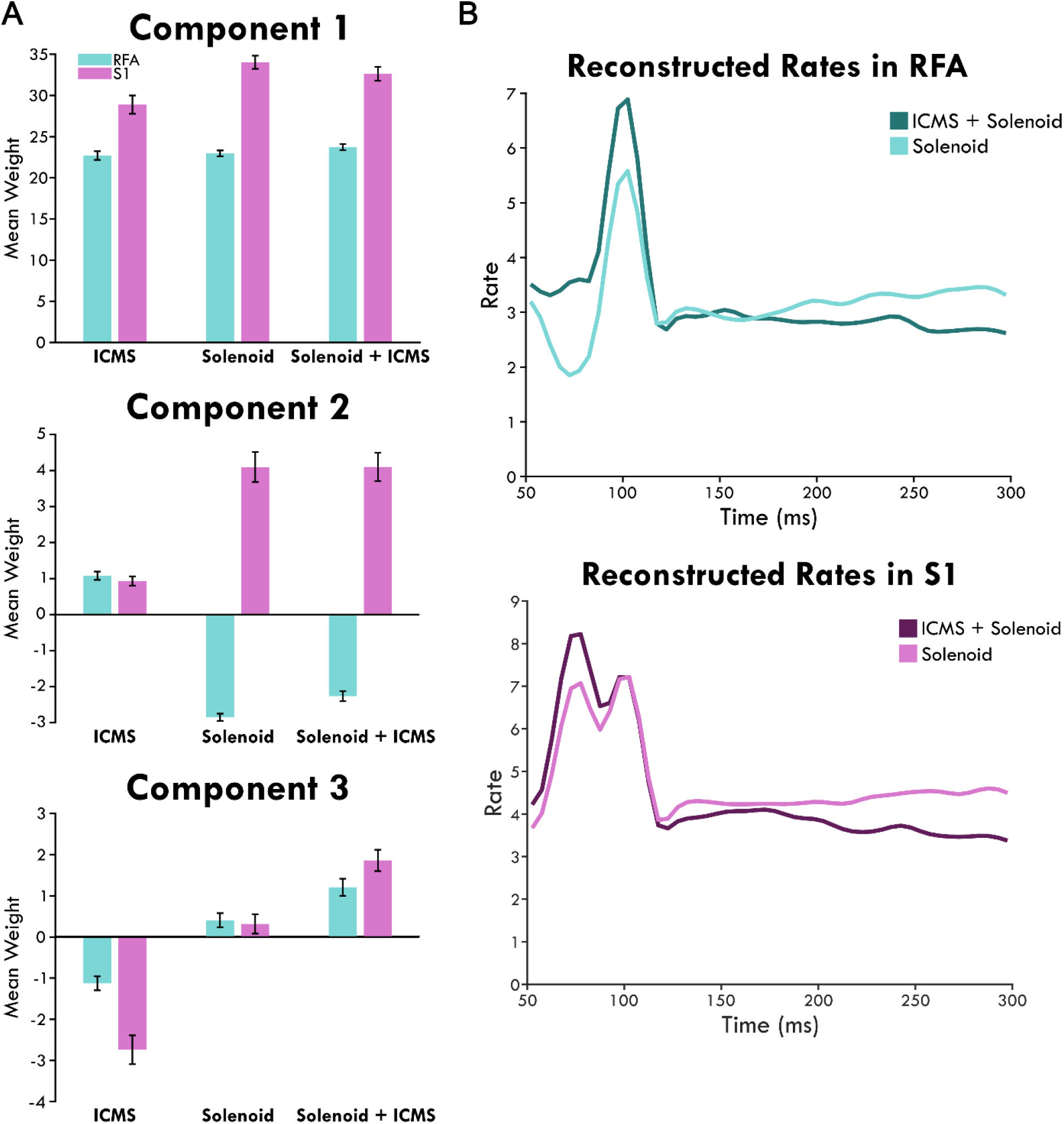
The mean weight of each independent component by area and stimulation type and their reconstructed weights. Panel A shows bars graphs with the mean independent components of trials sorted by stimulation type: ICMS, Solenoid, and ICMS + Solenoid. The bars show the mean explained by the top three independent components of each stimulation type in the rostral forelimb area (RFA) shown in blue and sensory cortex (S1) in pink. An inverse relationship is indicated by the negative values. The error bars show the standard error of the mean. Panel B shows the reconstructed rates for each stimulation type by area. The mean component scores were used to weight each component and reconstruct the average response in spiking to stimulation.

Using the mean component scores for each area, the rates of spiking activity can be reconstructed. Figure 6B shows the modeled rate for Solenoid and ICMS + Solenoid. There is a late peak at 100ms in RFA, 50ms after solenoid onset, in the rate of both trial types. The peak with ICMS + Solenoid is higher than with Solenoid alone and the dip before it disappears. The rates in S1 are bimodal with two peaks at 75 and 100ms. The rate for ICMS + Solenoid has a higher early peak compared to Solenoid trials but are otherwise similar.

## 4 Discussion

The primary purpose of this study was to probe the connections between somatosensory and premotor cortex after injury to the primary motor cortex by defining the somatosensory-evoked response in the average spike rate and modulating it through the application of a leading ICMS pulse. To assess the modulation of the population spiking rate relative to stimulation, we utilized independent components analysis, using the top three independent components of the data recorded in RFA and S1 for each trial type. These computed independent components were used to reliably describe the response in both areas separately and across stimulation types. Component 1 was time invariant and was related to a general increase in firing and did not provide information relative to stimulation events and was dropped from further analysis while both Components 2 and 3 had clear peaks that were related to increases in spiking activity. We found that there was a response to the solenoid strike in both cortical areas that varied in profile based on lesion volume and stimulation type. The results support the conclusion that recovery from CFA injury alters cortical processing of somatosensory responses, which may be involved in the restitution of sensorimotor integration.

### 4.1 Assessment of the lesion and its impact on cortical sensory responses

Motor information is received in the sensory cortex which can then modulate motor activity [31, 32]. The somatosensory-motor network is underpinned by direct corticocortical and corticospinal connections as well as subcortical structures such as corticostriatal and corticothalamic loops [33, 34, 35]. The crosstalk between sensorimotor areas allows for sensory predictions and feedback correction to fine tune the motor output [5, 36, 37]. Injury to the motor cortex can result in somatosensory deficits, further exacerbating the observed motor impairments [9]. These somatosensory deficits may be mediated by a loss of inhibitory input from primary motor cortex which is important for temporal coding [38]. In our study, variable lesion volumes and locations led to heterogeneous behavioral results, but, in general, rats that had larger lesions and greater cortical and subcortical involvement showed increased severity of injury. We expected concurrent modulation of somatosensory information with lesion volume since the cortico-cortical connections that mediate it were differentially impacted by injury. As a result, we found that lesion volume predicts trends in the spiking response of both cortical areas regardless of stimulation type. The extent of the lesion and the anatomic involvement of neural populations appears to have a significant effect on the sensory response such that any additional modulation by ICMS is obfuscated. Previous work has established the importance of lesion volume and location as predictors for the severity of motor impairment, but the impact on the somatosensory response as well solidifies its importance in sensorimotor processing over recovery [39].

### 4.2 Cortical representation of the sensory response after recovery

The premotor cortex is an important site for motor recovery after damage to the primary motor cortex. In addition to extant projections between S1 and the rodent premotor cortex, recovery is often coupled with the formation of de novo connections [14, 40]. Ischemic injury and the disruption to established connections initiates a cascade of compensatory processes including the upregulation of growth associated genes which can promote dendritic branching and synaptogenesis, supporting functional recovery [41, 42]. The effects of the molecular response extends beyond the infarct to other cortical areas [43]. While sparse connections exist between RFA and S1, we expect that as premotor cortex takes over function after injury to M1, connectivity to S1 will be strengthened. In uninjured animals, the premotor cortex shows increased activity in response to peripheral somatosensory stimulation; however, the process has been proposed to be mediated by primary motor cortex [22, 44]. After injury to CFA, there is a clear response to peripheral stimulation in the somatosensory cortex. Interestingly, we also observed a response 50 ms after the solenoid strike in RFA despite the damage to CFA, which hints at post-ischemic adaptations to restore sensorimotor integration. In S1, where we expect the primary processing of the solenoid stimulus to occur, this same peak exists in the average solenoid response; however, there is also an earlier peak in activity which is unique to S1, resulting in a bimodal response with peaks at 25 and 50 ms relative to solenoid strike. This early peak in S1 is more likely a traditional somatosensory response associated with somatotopic representation as it aligns with previously cited latencies [18] while the later peak 50 ms after solenoid onset may represent some coding of the sensory information with the variation in response profile dependent on the cortical layer and the nearby cell types since its presence on a channel basis is largely variable.

### 4.3 Evidence for the reorganization of somatosensory processing

Because both RFA and S1 have a similar response which is constrained to the same time point, there must be some factor involved in their synchronicity. The solenoid contact was set to last for 50 ms so the offset would correspond to the later peak in the response; however, the conduction time for a peripheral event such as solenoid offset would have to be nearly instantaneous [45]. Instead, it is more likely that the response is the result of central communication. Neuronal conduction between S1 and RFA was primarily found to be a function of the primary motor cortex under normal conditions and calculated to be on the order of 10 ms, so the longer latency response implies a disruption of normal pathways and subsequent adaptations [18, 22, 37]. Recovery of the somatosensory response in RFA could alternatively be mediated by thalamocortical projections as the 50 ms latency after solenoid onset does however resemble peak spiking latencies in the thalamus [46]. The synchronous peak might then be explained by this shared input which could be the product of post-injury reorganization. On the other hand, new projections between S1 and RFA as have been shown in previous study may have a longer latency without the mediation of primary motor cortex and could also coordinate the late somatosensory response [14]. As a result, the source of the somatosensory evoked peak in spiking within the premotor cortex—whether though direct projections from somatosensory cortex or a shared input—should be further investigated. The effect of lesion volume and area predicts an inverse relationship between Components 2 and 3 of trials in S1 regardless of trial type—with larger lesion volumes the mean weight of Component 2 goes down while Component 3 increases. This could represent a shift from the earlier to the later response particularly as integration of the premotor and somatosensory areas is no longer mediated by the damaged primary motor cortex, instead re-establishing mutual information between the areas after significant disruption. Other work, however, has shown that there is inhibition from primary motor cortex in S1 which is disrupted in injury and the peak increase with larger lesions may alternatively be due to disinhibition [38]. In either case, the significance of the later peak of the sensory response is emphasized after injury and, since it is shared by both RFA and S1, supports the plasticity of sensorimotor integration.

### 4.4 Modulation of the solenoid evoked sensory response

The ICMS pulse was used to probe the connectivity of the two cortical areas by determining if evoked activity in RFA influences the somatosensory response of S1. Analysis of the spiking activity after ICMS-only stimulation highlights a trend for an initial increase in activity which drops with time from the stimulus pulse as represented by Component 2. In RFA where the ICMS stimulation occurred, there was a clear peak in spiking after ICMS delivery. When the ICMS pulse preceded the solenoid stimulation, Component 3, corresponding to the peak 50ms after the solenoid response, was predicted to have a significantly greater mean weight than solenoid stimulation alone, meaning that on average ICMS facilitated spike firing at this timepoint. Our reconstructed rates further substantiate this by showing how ICMS + Solenoid trials have an elevated peak in RFA. ICMS delivered in RFA shifted the relative firing properties of the neurons within the immediate vicinity of the pulse and likely impacted the processing of somatosensory information. The influence of ICMS evoked activity in RFA on the somatosensory response in S1 is more difficult to disentangle. The mean weight of Component 3 is also predicted to be higher in S1 with ICMS + Solenoid stimulation, but this did not impact the reconstructed peak in mean spiking as expected. Furthermore, there was no effect of stimulation type on Component 2. Changes in the mean weight of Component 2 may be non-significant, but we see there is an increase in the reconstructed rate at the corresponding early part of the response in both areas with the addition of ICMS despite this. It is important to note that there are differences in the component shapes between the stimulation types which may account for some of these discrepancies. The shape of the independent components for ICMS + Solenoid resembles those for the Solenoid trials, the exception being Component 2 which shifts to include a negative going portion. The result of this is that Component 2 also affects the later response that corresponds to Component 3. The goal of ICA was to isolate independent features of the response; thus, the change in the Component 2 to include both an early and late profile after the ICMS pulse suggests that the two response patterns become more related; however, this may also be an artifact of the component fit between the two areas. Regardless, it is important to note that the components are derived to best capture the data so a change suggests that ICMS has sufficiently altered the shape of the sensory response to require new descriptors. We found that there is a definitive shift that occurs with the addition of ICMS represented in the distribution of channel-averaged activity along the component axes. The underlying rates vary with the changes in the representative weights; many increasing their firing rate with some decreases as well. Altogether, there is a clear disruption of normal sensory processing with ICMS + Solenoid in S1 that can be attributed to the influence of RFA activity in recovery. The independent components may not be a perfect fit for the mean data but do better at capturing trial to trial variations; therefore, the subtle changes in mean weights and reconstructed rates are only hinting at the diverse population of responses that underly the modulation of sensory processing.

### 4.5 Inter-channel variability of spiking activity and cortical layer dependence

This study combined data from different animals, cortical layers, lesion and somatosensory stimulus delivery conditions which contributed to a large amount of heterogeneity in the somatosensory response that was obfuscated by the channel averaged mean. The confidence intervals of the peri-event time histograms had spreads that surpassed the baseline spiking threshold as shown in Figure 3, hinting at the underlying channel variability. Each channel’s depth in the cortex depended on the relative depth of the array and the location of the channel on the electrode shank. As a result, the channels measured the relative contribution of neural populations spanning the cortical column; however, projections between sensorimotor cortical areas are layer specific and likely influence the response of local neural populations differently [38, 47]. This study was limited in the ability to discern the location of the electrode site and we chose not to focus on analyzing the averaged response by the cortical layer. However, further studies should examine the contribution of layer-specific somatosensory processing within the cortical column as it could provide insight into participating pathways and cell types.

### 4.6 Summary

Overall, we found that there was a response in spiking activity in both RFA and S1 even after lesioning of CFA. Part of the somatosensory response is shared between the two cortical areas and is altered as a function of injury. Component analysis suggests shifts in the response patterns of S1 as a result of evoked activity in RFA, implying a functional relationship between RFA and S1 which may be involved in somatosensory processing. Together, these results highlight the direct recruitment of premotor cortex in somatosensory processing and its modulatory effects on somatosensory cortex. The relationship between the RFA and S1 is further evidence for the re-establishment of sensorimotor integration after injury.

## 6 Author Contribution

MDM and DJG designed the experiment with input from RJN and PH. MDM, PH, CT, SD, JB carried out the experiment. PH and MDM analyzed the data. PH interpreted the results, created figures, and wrote up the manuscript with the help of MDM. RJN and DJG edited and guided manuscript development.

## 8 Acknowledgements

Thanks to Jasmine Deng for her surgical assistance.

