## Supplementary Figures for "Post-Ischemic Reorganization of Sensory Responses in Cerebral Cortex"

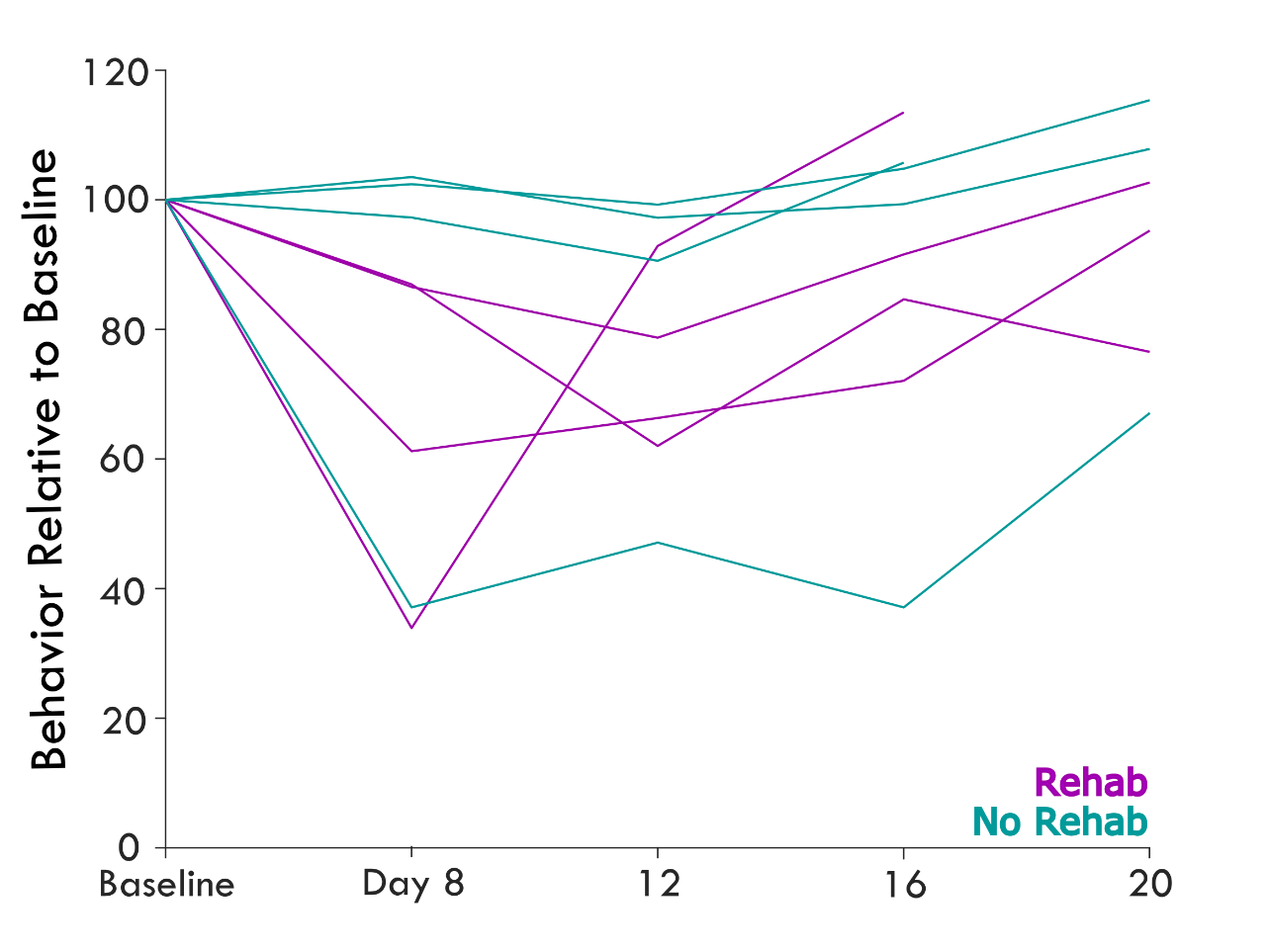
**Supplementary Figure 1. Single pellet reach task success for individual rats by treatment.** Each line is the percent success in behavioral performance across the experiment relative to baseline which has been standardized to one hundred percent. The purple color shows rats belonging to the group which received rehabilitative training and the teal color shows those not receiving any intervention after injury.

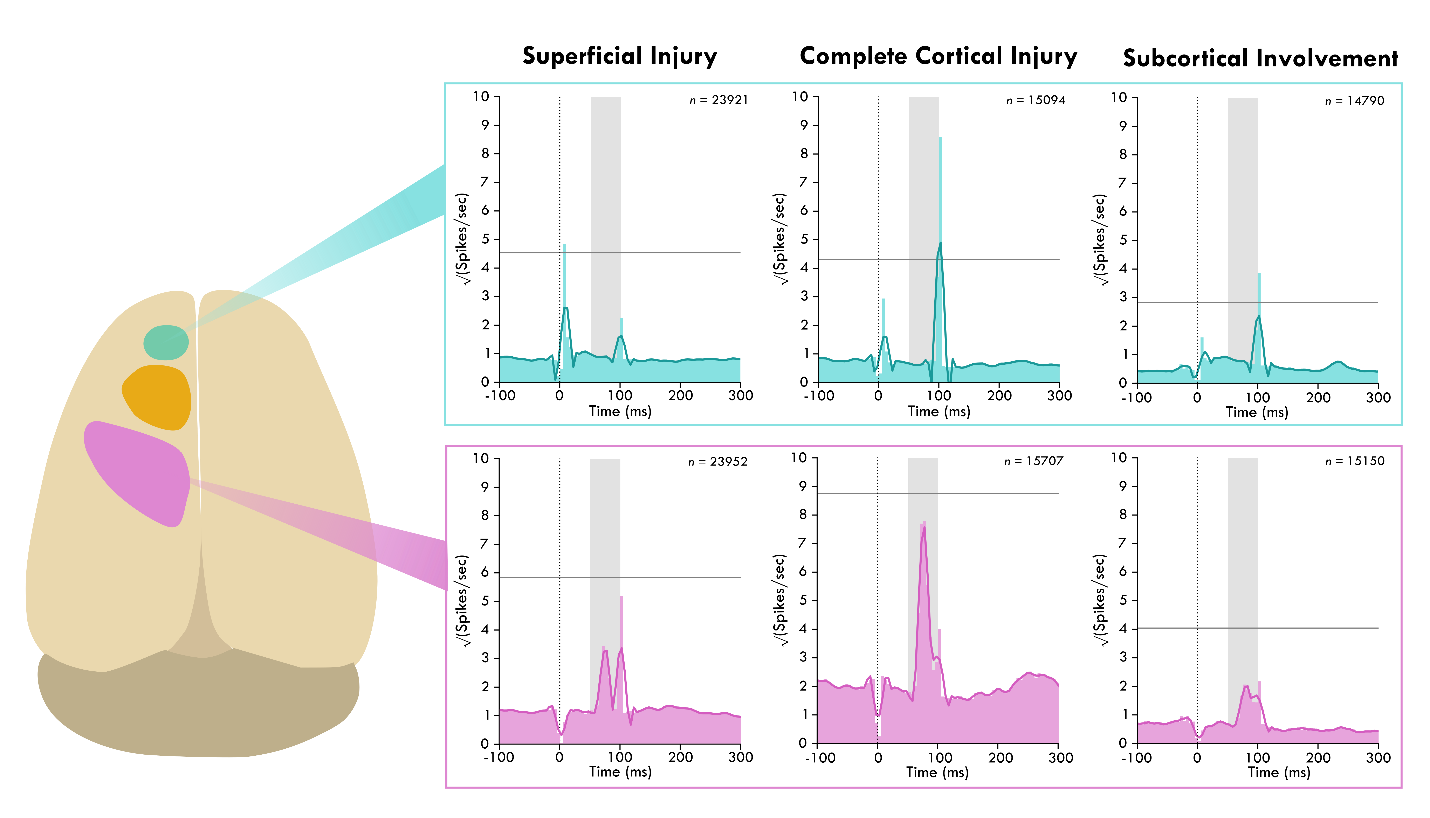
**Supplementary Figure 2.** **Average spiking response of different injury categories to solenoid strike with a preceding ICMS pulse.** The schematic shows the cortical areas and the average spiking activity for lesion type groups categorized by depth: *Superficial Injury*, where the lesion only impacts the upper cortical layers (MM-S1, MM-U1, and MM-U2); *Complete Cortical Injury*, where the lesion impacts all cortical layers (MM-V1 and MM-W1); *Subcortical Involvement*, where the lesion impacts all cortical layers and impinges on subcortical territory (MM-T1 and MM-T2). Each peri-event time histogram shows the square root of the spikes in each time bin and the smoothed mean as a darker overlaid line. ICMS delivery is represented as the vertical dotted line and solenoid activity by the shaded gray area with an onset and offset of 50 and 100ms respectively.  The gray horizontal line is one standard deviation above the average pre-stimulus activity.

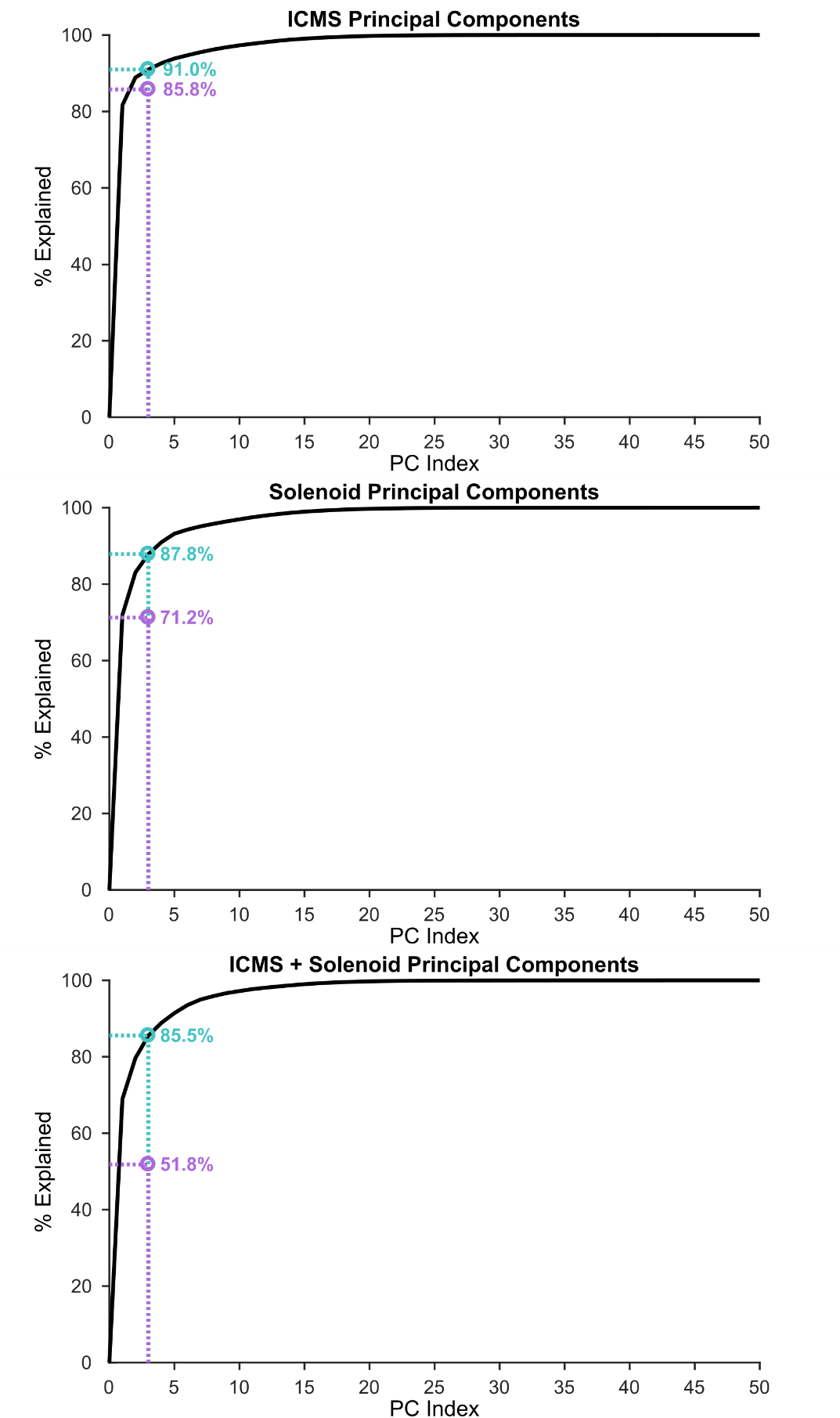
**Supplementary Figure 3.** **The percent of the signal explained by the top three components in each stimulation type.** The number of principal components is shown on the bottom where the line shows how the signal is accounted for in the percent explained on the y-axis with the addition of more principal components. The independent components are derived from the principal components, but the reconstructed axes are rotated to be statistically independent of one another. The purple point marks the percent of the data explained with the newly reconstructed axes.

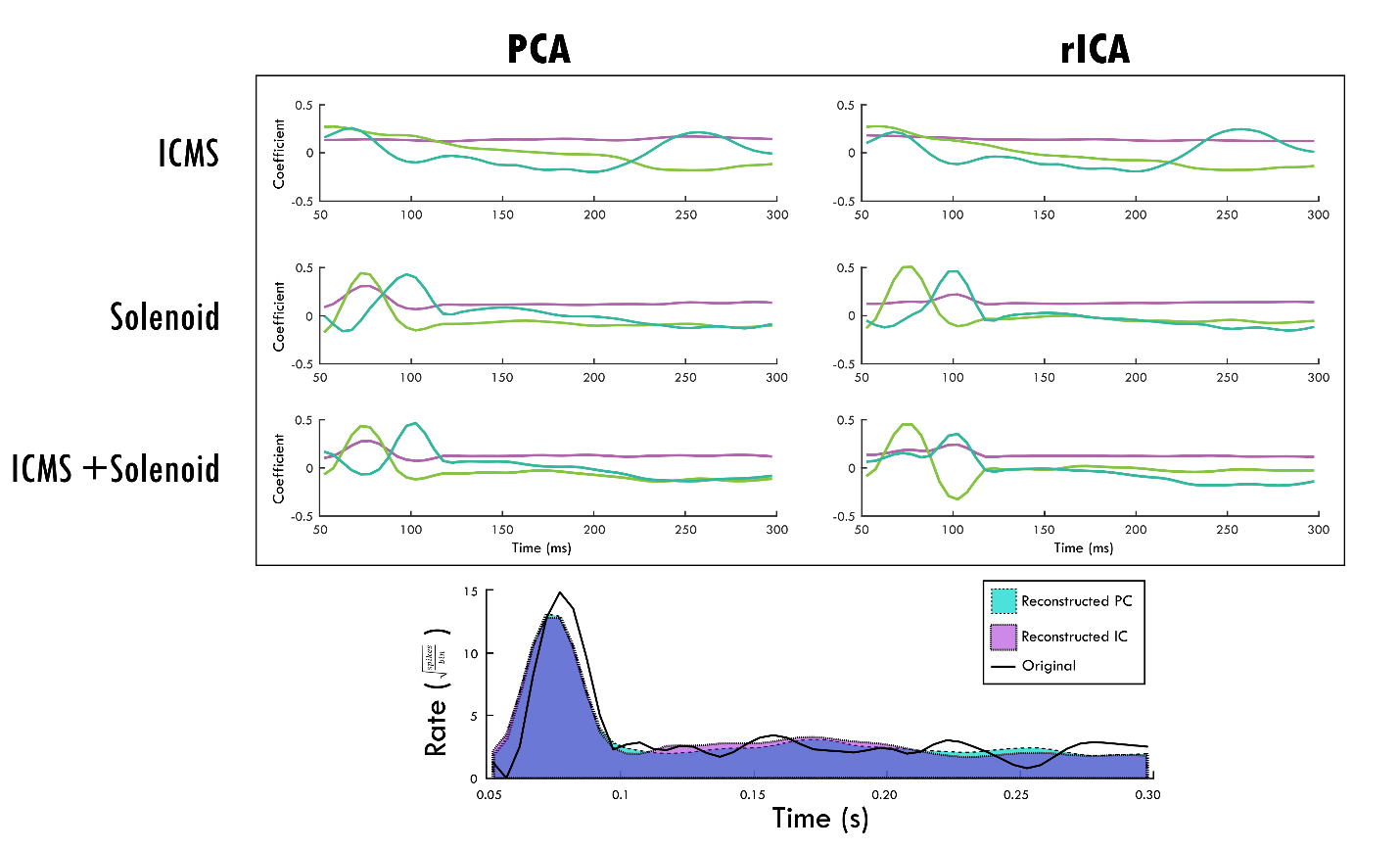
**Supplementary Figure 4. The difference in principal components and reconstructed independent components.** The top-3 components of both methodologies are shown for each stimulation type. The color order goes from purple, green, to blue. The bottom panel shows an example trial (solid black line) and its reconstruction by each methodology as a transparency.
